# Mitotic centromeres organize in repeat-subunits with specialized histone landscape and multi-domain architecture as revealed by expansion microscopy

**DOI:** 10.64898/2026.10.30.763054

**Authors:** Gokilavani Thangavel, Sylvia Erhardt

## Abstract

Centromeres ensure faithful chromosome segregation during cell division, yet their underlying architecture and spatial arrangement of centromeric proteins remain incompletely resolved. Here, we apply Ten-fold Robust Expansion Microscopy (TREx) to human diploid hTERT RPE-1 cells to visualize mitotic centromeres at nanoscale resolution. We find that the centromeric histone variant CENP-A and the kinetochore protein KNL1 form discontinuous, repeating substructures. We further resolve centromeric chromatin into four spatially distinct domains: an inner kinetochore domain marked by CENP-A, a mantle domain marked by RAD21, an outer core domain marked by CENP-B, and a distinct, RAD21-associated inner centromere domain. Core and pericentromeric regions share a specialized chromatin landscape, extending the concept of centrochromatin, with enrichment of H3S10ph and H3K9me3, depletion of H3K27me3, and low-level enrichment of the euchromatic marks H3K4me3 and H3K36me3. Together, we uncover a discontinuous repeat-subunit organization and multidomain architecture of the human centromere that combines euchromatic and heterochromatic modifications.

## Introduction

Centromeres are essential chromosomal loci that direct kinetochore assembly and thereby ensure accurate chromosome segregation during cell division. Despite decades of investigation, their architecture and protein composition remain incompletely understood. This is largely because centromeric DNA is highly repetitive and compact. These features have limited both, short read sequencing, which cannot precisely map repeat arrays, and conventional light microscopy, which cannot resolve structures below about 250nm. In humans, centromere identity is specified epigenetically by the histone H3 variant CENP-A, one of the centromere proteins first identified with autoimmune sera and which was reported organized into distinct foci (Earnshaw & Rothfield, 1985). By conventional microscopy, each sister centromere appears as a single diffraction limited focus of CENP-A. However, several lines of evidence indicate that this focus contains considerable substructures.

In 1989, the plant *Haemanthus* was reported to display a distinct double-dot pattern for centromeres in each chromatid, which the authors termed a “bipartite structure” (Mole- Bajer et al., 1990). Soon after, analysis of fragmented kinetochores in several species led to the “repeat subunit model”, in which the kinetochore is built from multiped repeat, repeated functional units (Zinkowski et al., 1991). Consistent with this model, stretched chromatin fibers revealed that CENP-A nucleosomes are not contiguous but are arranged in blocks interspersed with blocks of canonical H3 nucleosomes (Blower et al., 2002). And more recently, expansion microscopy confirmed bipartite centromeres in human, mouse and chicken cells and extended this organization to the kinetochore (Sacristan et al., 2024). In agreement with this model, a recent cryo-electron tomography study revealed multiple CENP- A chromatin regions separated by CENP-A-depleted regions on a single chromatid of humans (Kixmoeller et al., 2025).

Telomere-to-telomere assemblies of human centromeres now allow the questions of precise CENP-A occupancy to be addressed at the sequence level. CENP-A-bound DNA regions coincide with a locally hypomethylated stretch of alpha satellite DNA, called the centromere dip region (CDR) (Altemose et al., 2022; Gershman et al., 2022), and many centromeres contain more than one dip region or the dip region is slit into several sub-dip regions (Gao et al., 2026). Recent long-read sequencing studies revealed that there are between one and ten CENP-A subdomains within the region that binds CENP-A (Gao et al., 2026; Xu et al., 2026). Together, these studies show that the CENP-A domain is more complex and may organize in several subdomains that is in need for a more detailed structural analysis.

In humans, CENP-A-containing chromatin which mediates microtubule attachment, constitutes only a very small fraction of the total centromeric alpha satellite array region (Marshall et al., 2008). This raises fundamental questions about the chromatin composition and higher-order organization of the centromeric chromatin domain as a whole. Accumulating evidence suggests that human centromeric chromatin is not uniform but instead is organized into multiple functionally distinct domains. We recently reviewed evidence supporting the existence of at least four functionally distinct domains (Corless et al., 2025): The inner kinetochore domain, composed of CENP-A containing chromatin, forms the first layer of centromeric chromatin and mediates kinetochore assembly and microtubule attachment. Beneath this lies the mantle domain, which remains poorly characterized. Recent observations suggest that cohesin and condensin complexes localize beneath CENP-A, potentially contributing to the organization of the CENP-A domain (Sacristan et al., 2024; Sen Gupta et al., 2023). The outer core domain, which comprises the majority of centromeric chromatin, is associated with CENP-B binding to the 17-bp CENP-B box within α-satellite centromeric DNA (Gamba & Fachinetti, 2020). This domain may provide structural support. The inner centromere domain has been implicated in sister chromatid cohesion and contain proteins of the chromosomal passenger complex and structural maintenance of chromosomes complexes (Gireesh et al., 2025; Hengeveld et al., 2017). Collectively, these observations point to a multidomain architecture in which only the CENP-A-containing inner kinetochore domain constitutes the core centromere, whereas the surrounding domains form three-dimensional distinct subdomains of the pericentromere, which is required for the establishment and maintenance of core centromeres.

Post-translational modifications of histones influence chromatin structure and function. The chromatin interspersed with CENP-A has a distinctive modification pattern, termed centrochromatin. Studies on extended chromatin fibers detected that it carries H3K4me2 but lacks the H3K9 methylation that marks the flanking pericentric heterochromatin (Sullivan & Karpen, 2004). Further marks have since been described within this domain, including H3K36me2 (Bergmann et al., 2011) and H4K20me1 (Hori et al., 2014). Combined with DNA hypomethylation at the CDR, these marks indicate that the core centromere is epigenetically distinct from its surroundings. However, the spatial distribution and organization of histone modifications within human centromeric chromatin has not been examined using high resolution imaging and it is unknown how these marks relate to CENP-A subdomains in three dimensions.

Super-resolution microscopy now offered a way to resolve the three-dimensional architecture of mitotic centromeres and the molecular composition of centromeric chromatin. Expansion microscopy physically enlarges biological specimens, increasing the distance between fluorophores and thereby enabling nanoscale resolution using conventional imaging platforms (Damstra et al., 2023). Past studies have employed 4.5x expansion microscopy to study centromere architecture (Di Tommaso et al., 2023; Sacristan et al., 2024). By combining Ten-fold Robust Expansion Microscopy (TREx, Damstra et al., 2023) with immunofluorescence for centromeric proteins and histone modifications, this study aims to resolve the number of sub-domains underlying the CENP-A occupied core centromeric region, as well as characterize the multidomain organization and histone modification landscape of entire centromeric chromatin. We find that the CENP-A domain of a single mitotic centromere comprises of 1-9 distinct subdomains, rather than thesone or two foci previously described. This number is consistent with the multiple hypomethylated dips detected at the CENP-A- binding region by long-read sequencing.

## Results

### Visualizing centromeres using expansion microscopy

To determine the number of CENP-A domains within a single centromere, we established Ten-fold Robust Expansion Microscopy (TREx; (Damstra et al., 2023) in human cell line hTERT RPE-1. TREx was designed for ten-fold expansion, but we obtained an expansion factor of approximately seven-fold (expansion factor = diameter of the gel after expansion (cm) / diameter of the gel before expansion (cm)). This is likely due to the batch variability of sodium acrylate quality, a known issue with the method (Damstra et al., 2023). Combined with Airyscan super resolution imaging (with a lateral resolution of 120-140nm), we achieved a resolution of at least 17-20 nm, higher than most fluorescence-based imaging resolution of human centromere studies reported so far. To preserve the native three-dimensional architecture of metaphase centromeres we first performed TREx on cells grown on coverslips without any drug treatment and screened for cells in metaphase. To estimate the number of CENP-A domains, we generated intensity profiles for individual centromere and counted peaks (Fig. S1). Centromeres contained between one and eight CENP-A peaks, and the largest fraction (42.6%; n = 423 chromatids from 7 cells) showed two distinct peaks (Fig. 1a,b). This is consistent with the bipartite organization previously observed by 4.5-fold expansion microscopy (Sacristan et al., 2024).

**Figure 1.**
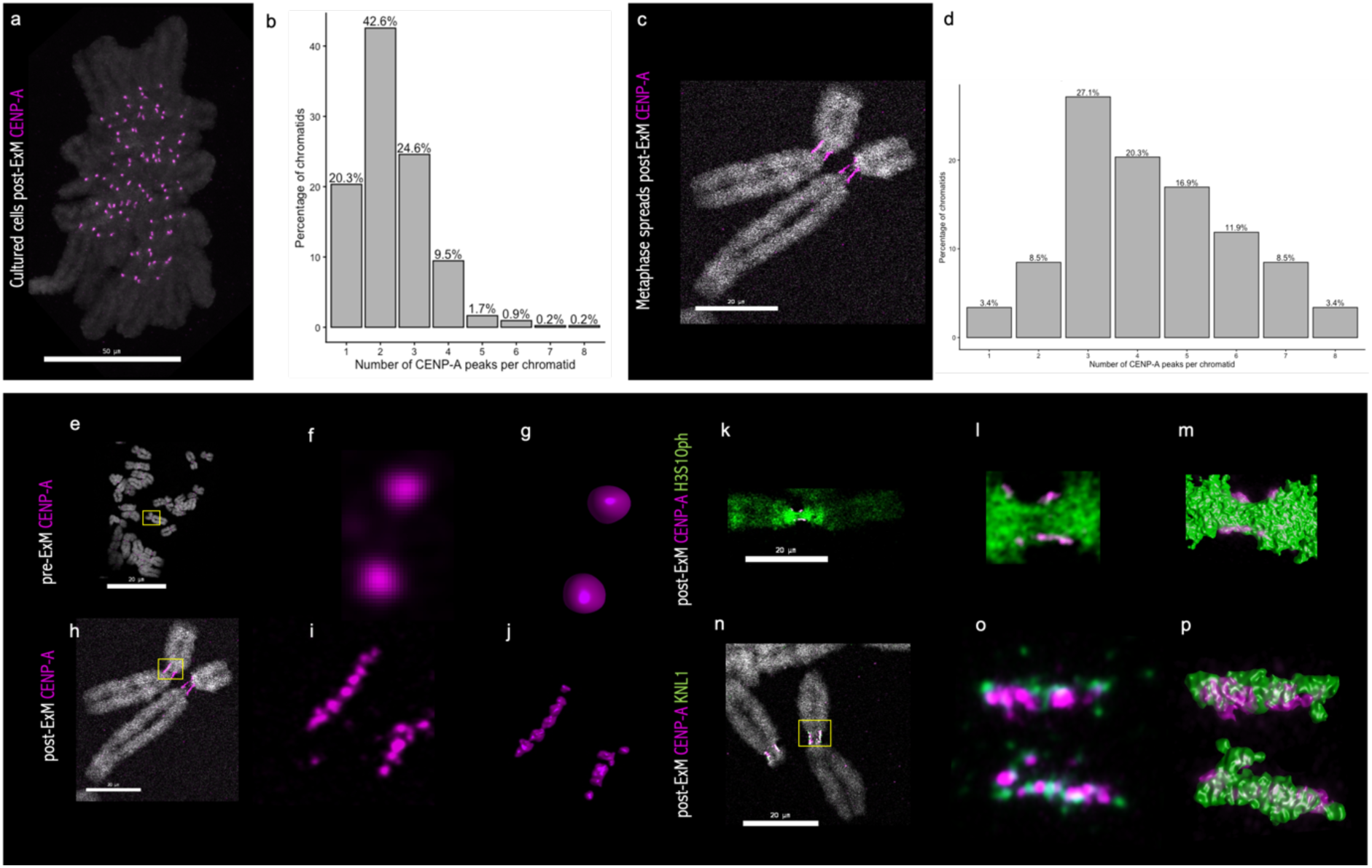
Visualizing the three-dimensional architecture of centromere and kinetochore proteins using expansion microscopy. **a**, CENP-A staining (magenta) on post-expanded metaphase cells cultured on coverslips. **b,** Distribution of CENP-A foci per individual chromatid in native conformation, the number of chromatids analyzed were 423 **c,** CENP-A staining (magenta) on post-expanded metaphase chromosome spreads **d,** Distribution of CENP-A foci per individual chromatid in metaphase spreads, the number of chromatids analyzed were 59. **e–g,** CENP-A staining (magenta) on pre-expanded metaphase spreads appears as single foci. **h–j,** CENP-A staining on post-expanded metaphase spreads resolves into multiple discontinuous subunits linearly arranged along the primary constriction. **k–m,** CENP-A (magenta) and H3S10ph (green) staining reveals H3S10ph signals between discontinuous CENP-A subunits**. n–p,** CENP-A (magenta) and KNL1 (green) staining reveals multiple discontinuous subunits for both proteins. Scale bar, 20 μm.

In intact metaphase cells, centromeres are compact and randomly oriented relative to the imaging plane. This means that adjacent subdomains may overlap, especially along the optical axis, where resolution is lower. To increase the spatial separation between subdomains, we applied TREx to metaphase spreads from colcemid-treated cells obtained by hypotonic swelling followed by cytocentrifugation (Hsu & Pomerat, 1953). Cytocentrifugation flattens and partially extends mitotic chromosomes. It disturbs their native three-dimensional conformation, but the linear order of chromatin along the centromere should be preserved, so subdomains that overlap in intact cells can be resolved, enabling us to capture the precise underlying CENP-A domain counts which is otherwise difficult to resolve in native conformation. In spread cells, CENP-A appears as multiple discontinuous foci arranged linearly along the primary constriction (Fig. 1c,h–p, Fig. S1). Individual sister chromatids contained between one to eight CENP-A foci, with most ranging between three and five (Fig. 1c,d) (median is 4; n = 59 chromatids from 28 cells). This range is consistent with the CENP-A sub- domain count based on recent long-read sequencing and DNA methylation profiling (Gao et al., 2026; Xu et al., 2026).

Because chromosome spreading and expansion microscopy are techniques that apply forces on the biological specimens, we considered whether these multiple substructures arise from fragmentation of the chromatin fibers. We reasoned that if the gaps between CENP-A foci reflect breaks, they should be devoid of chromatin. We therefore costained CENP-A with the H3-specific phosphorylation antibody H3S10ph. In mitosis, this phosphorylation of H3 is present on condensed chromatin starting from pericentromeric heterochromatin (Hendzel et al., 1997). H3S10ph signals was consistently detected interspersed between the multiple CENP-A foci (Fig. 1k–m). Thus, this strongly indicates that a single CENP-A locus is indeed composed of multiple substructures interspersed with H3-containing chromatin visible at nanoscale resolution. This matches the interspersed organization first seen on stretched chromatin fibres (Blower et al., 2002). We next examined the outer kinetochore protein KNL1 with TREx. KNL1 also formed multiple discontinuous foci, suggesting that the sub-domain organization of CENP-A chromatin expands to the outer kinetochore as well (Fig. 1n–p).

### Visualizing the spatial organization of centromeric chromatin domains

To test the multi-domain model of centromeric chromatin (Corless et al., 2025), we applied TREx to visualize markers of each proposed domains: CENP-A of the inner kinetochore domain, the cohesin complex component RAD21 for the mantle domain and inner centromere domain , and CENP-B for the outer core domain. CENP-A formed linear arrays on the poleward surface of the centromeric chromatin and occupied only a fraction of the primary constriction. This is consistent with a spatially restricted inner kinetochore domain within the larger centromeric region (Fig. 2a–d).

**Figure 2.**
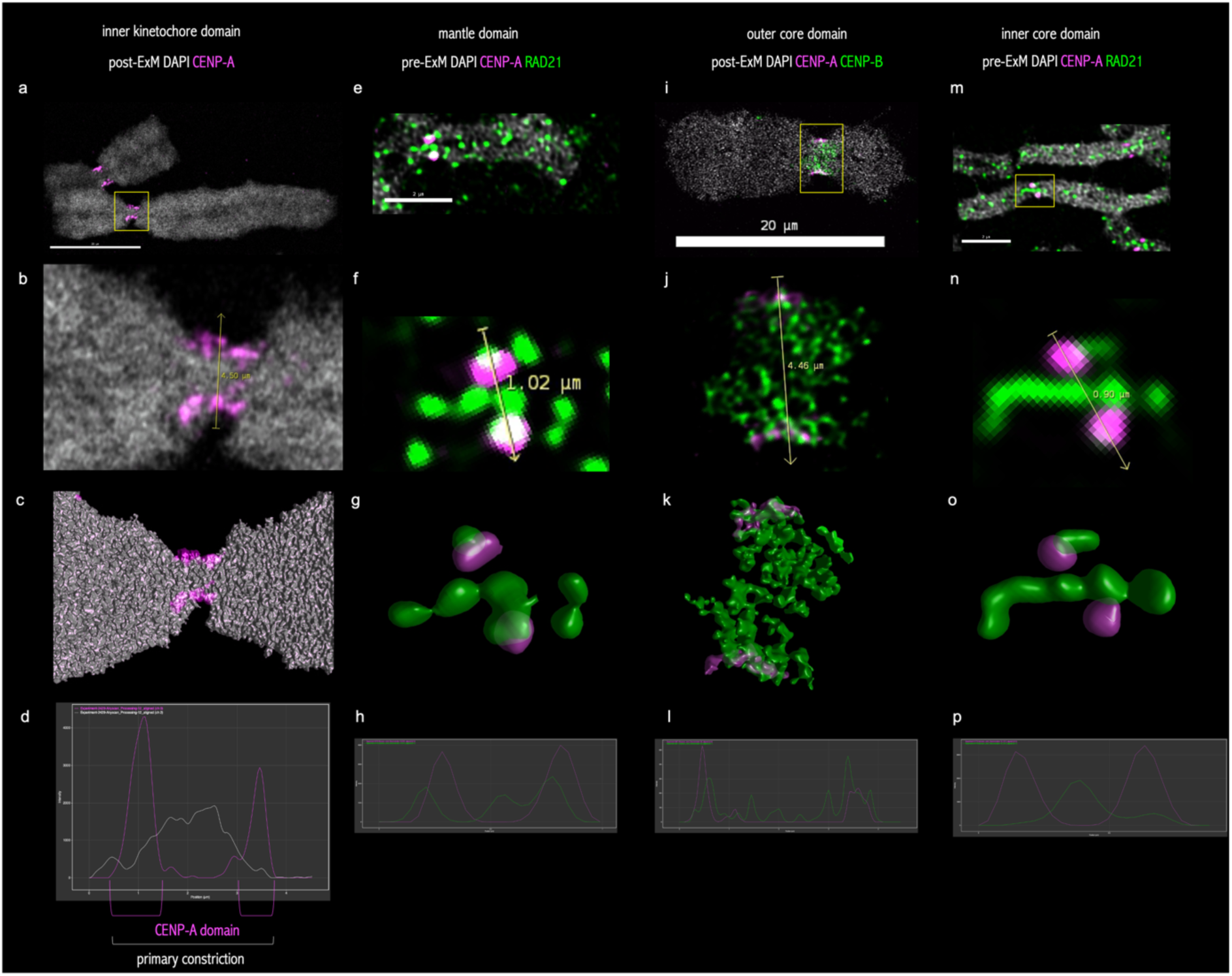
Visualizing the multi-domain architecture of the centromere using expansion microscopy. **a–d**, CENP-A (magenta) and DNA (grey). CENP-A occupies only a fraction of the primary constriction, defining the inner kinetochore domain**. e–h,** CENP-A (magenta) and RAD21 (green), showing the co-localization of RAD21 with CENP-A, indicative of the mantle domain**. i–l,** CENP-A (magenta) and CENP-B (green), showing the distribution of CENP-B throughout the primary constriction. CENP-B occupies the outer core domain and extends across the primary constriction, contributing to all four centromere domains. **m–p,** CENP-A (magenta) and RAD21 (green), showing RAD21 localization along the chromosome axis, defining the inner-core domain. Scale bar, 20 μm.

While examining the mantle domain, we realized that RAD21 signal was too low for applying TREx and therefore analyzed pre-expanded mitotic centromeres. While consistently present in a continuous linear signal between sister chromatids, which indicated that there is a retention of centromeric cohesin after the prophase pathway removes cohesin from chromosome arms, and it supports the existence of an inner centromere domain. (Fig. 2e–h, m–p). By contrast, RAD21 signal adjacent to CENP-A, which would mark the proposed mantle domain, was variable. It was observed in only 16% of mitotic chromosomes (n = 1140 chromatids from 89 cells) (Fig. 2e–h, Fig. S2g). At the resolution of unexpanded signals, we cannot determine whether this signal lies beneath CENP-A as the mantel model predicts.

CENP-B, which binds the 17-bp CENP-B box in α-satellite DNA, stained the entire primary constriction. This is consistent with α-satellite chromatin forming the bulk of the centromere region as predicted for the outer core domain (Fig. 2i–l). CENP-B signals also overlapped with CENP-A, consistent with CENP-B boxes being present within the CENP-A- bound α-satellite arrays and its occupancy stretches along the whole centromeres. The outer core domain extended into the inner kinetochore and mantle domains. Interestingly, on some chromosomes CENP-B was enriched on one side of the primary constriction (Fig. S2a–f). This may reflect an asymmetric CENP-B box density distribution or an uneven organization of the underlying α-satellite DNA itself. Finally, RAD21 was consistently detected between sister chromatids, confirming the presence of a distinct inner centromere domain (Fig. 2m-p).

### Visualizing the histone mark landscape of centromeric chromatin

To characterize the chromatin environment surrounding human centromeres, we examined histone modifications associated with transcriptionally active chromatin (H3K4me3 and H3K36me3), constitutive heterochromatin (H3K9me3), facultative heterochromatin (H3K27me3), and mitotic chromosome condensation (H3S10ph). CENP-A served as a marker of centromeric chromatin in all experiments. We performed TREx on interphase cells cultured on coverslips and metaphase chromosome spreads.

In interphase cells, CENP-A foci were frequently observed within DAPI-dense domains, consistent with the clustering of centromeric regions in heterochromatin-rich nuclear compartments. In late G2 cells, DAPI-dense regions showed strong H3S10ph signal, followed by prominent but weaker H3K9me3, whereas H3K27me3 signal was comparatively depleted (Fig. 3a,e,i). H3K4me3 and H3K36me3 were present at low but detectable levels within these domains (Fig. 3m,q). Only a small number of H3K4me3 or H3K36me3 positive foci were observed in close proximity to or overlapping with CENP-A foci.

**Figure 3.**
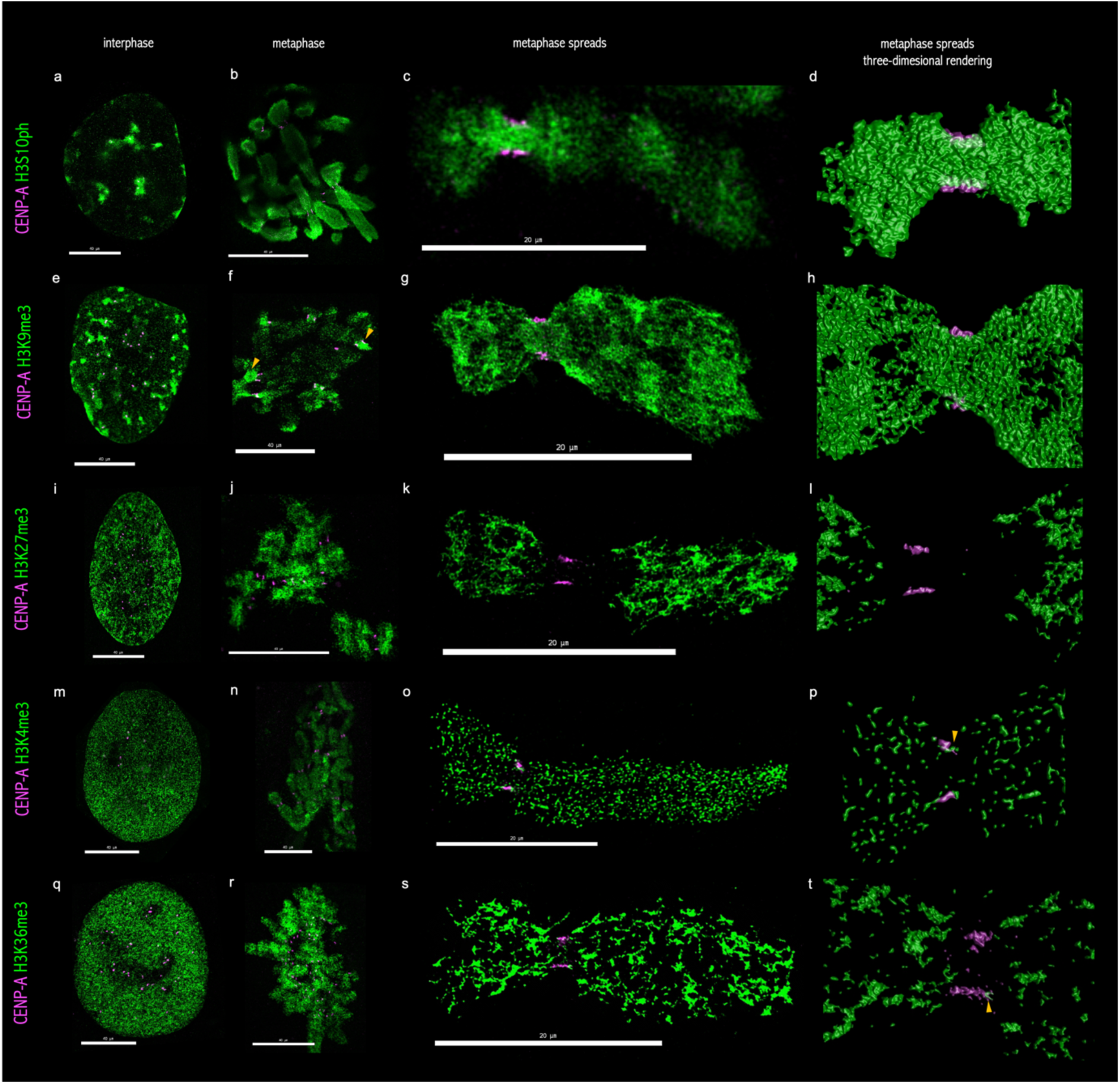
Visualizing the histone landscape of core and pericentromeric chromatin using expansion microscopy. **a,e,i,m,q**, Interphase images of adherent cells. **b,f,j,n,r,** Metaphase images of adherent cells. **c,g,k,o,s,** Metaphase images from chromosome spreads. **d,h,l,p,t,** Three-dimensional renderings of the corresponding metaphase spreads. **a–d,** CENP-A (magenta) and H3S10ph (green), showing strong enrichment of H3S10ph at centromeric regions**. e–h,** CENP-A (magenta) and H3K9me3 (green), showing enrichment of H3K9me3 at centromeric regions and banding pattern along chromosome arms. **i–l,** CENP-A (magenta) and H3K27me3 (green), showing depletion of H3K27me3 from centromeric regions and banding pattern along chromosome arms. **m–p,** CENP-A (magenta) and H3K4me3 (green), showing low enrichment of H3K4me3 at centromeric regions. **q–t,** CENP-A (magenta) and H3K36me3 (green), showing low enrichment of H3K36me3 at centromeric regions. Scale bars: 40 μm for **a,b,e,f,i,j,m,n,q,r** and 20 μm for **c,d,g,h,k,l,o,p,s,t.**

In metaphase chromosome spreads, we divided centromeric region into three regions: the first is the CENP-A-containing region, predominantly composed of core centromeric chromatin but potentially also including some proximal pericentromeric chromatin. The second is the inter-sister pericentromeric region, corresponding to the chromatin between the CENP-A signals of sister chromatids. The third is the arm-proximal pericentromeric region, extending towards the chromosome arms. Within the CENP-A containing region, H3S10ph showed the highest degree of spatial overlap with CENP-A, with a median colocalization value of 69% (Fig. 3b–d, Fig. S3). CENP-A itself lacks the H3 serine 10 residue, so this overlap reflects H3S10ph-marked H3 nucleosomes interspersed with CENP-A nucleosomes below the resolution limit. H3K9me3 also showed a substantial overlap, with a median colocalization of 27% (Fig. 3f–h, Fig. S3). In contrast, H3K4me3 and H3K36me3 showed minimal overlap, with median colocalization values of 0.2% and 0.5%, respectively (Fig. 3n–p,r–t, Fig. S3), and H3K27me3 was absent from this region, with a median colocalization of 0% (Fig. 3j–l, Fig. S3).

The inter-sister and arm-proximal pericentromeric regions showed similar histone- modification profiles. Both regions were enriched for H3S10ph and H3K9me3, whereas H3K27me3 was consistently depleted (Fig. 3c,d,g,h,k,l). On several chromosomes, H3K9me3 signal differed between the two arm-proximal pericentromeric regions, suggesting heterogeneity in the distribution of constitutive heterochromatin between chromosome arms (Fig. 3f). H3K4me3 and H3K36me3 were detected at low levels in both pericentromeric regions (Fig. 3o,p,s,t). Beyond the centromeric region, H3K9me3 and H3K27me3 formed distinct banding patterns along metaphase chromosomes, consistent with their localization to discrete chromatin domains (Fig. 3g,k). Together, these observations indicate that mitotic centromeric and pericentromeric chromatin are dominated by H3S10ph and H3K9me3, containing only low levels of H3K4me3 and H3K36me3, and lacks H3K27me3.

## Discussion

### Repeat subunit model of centromeres and kinetochores

Our CENP-A and KNL1 observations are consistent with the repeat subunit model of centromere-kinetochore complex, wherein a basic structure forms the basis of centromere organization and is repeated multiple times to form the complete centromere. This model was initially proposed by Zinkowski et al. (1991) based on cytological observations of stretched and fragmented metaphase chromosomes (Zinkowski et al., 1991). A recent cryo-electron tomography (cryo-ET) reached a similar conclusion: a single centromere locus contains multiple CENP-A occupied regions separated by CENP-A-depleted chromatin regions (Kixmoeller et al., 2025).

Sequence-level evidence points in the same direction. Because centromeric DNA is highly repetitive, short reads cannot be mapped unambiguously to it, and subdomain organization was therefore not visible in short-read sequencing data. Long-read sequencing and telomere-to-telomere assemblies have overcome this limitation. In these data, the CENP- A-bound region coincides with a locally hypomethylated stretch of α-satellite DNA, the centromere dip region (CDR) (Altemose et al., 2022; Gershman et al., 2022). Using Oxford Nanopore data from the HG002 genome, Xu et al. (2026) reported a mean of 5.5 sub-CDRs per centromere (range 2–9). DiMeLo-seq mapping of CENP-A in the same study gave similar numbers. In a population-scale analysis of 2,110 centromere assemblies, Gao et al. (2026) found between one and ten sub-CDRs per centromere, most commonly three to five.

The distribution from these sequencing approaches matches our microscopy-based observations of one to eight CENP-A foci per sister chromatid, with three to five domains being the most common. However, the comparison has some limits: Sequencing measures DNA methylation or CENP-A occupancy along linear sequence, averaged over many cells, whereas our imaging counts three-dimensional foci in individual chromosomes. Whether each CENP-A focus corresponds to a single sub-CDR remains to be tested. Nevertheless, expansion microscopy, cryo-electron tomography and long-read sequencing point to the same conclusion that the human CENP-A domain is not a single continuous block, but consists of multiple subdomains interspersed with H3 chromatin.

### Reconciling the repeat subunit and the bipartite model

An alternative model of centromere organization is the bipartite model of centromeres, initially proposed in 1990 in the plant *Haemanthus* (Mole-Bajer et al., 1990) and recently extended to human, mouse, and chicken centromeres (Sacristan et al., 2024). Using 4.5x expansion microscopy for centromeric proteins, the authors report mostly two subdomains per centromere. Together with staining for microtubule bundles and Capture-C, their data suggest each centromeres function as two units each interacting with its adjacent centromeres.

Our study shows that those two models are not mutually exclusive. In intact metaphase chromosomes imaged by TREx, we also most often observed two CENP-A peaks per centromere. Only in spread chromosomes, where subdomains are physically separated, did we resolve a higher number of CENP-A foci. This indicates that the bipartite pattern maybe a higher order organization consistent of several smaller subdomains of CENP-A. In terms of structure, microscopic observations (Kixmoeller et al., 2025) as well as long-read sequencing approaches (Gao et al., 2026; Xu et al., 2026), strongly indicate the number of CENP-A subdomains varies from one up to ten, supporting the repeat-subunit model of human centromeres, while functionally, the centromere might be bipartite that bind distinct microtubule bundles.

### A conserved basic unit across centromere types?

Centromeres across eukaryotes differ greatly in size and organization ranging from the 125bp point centromeres of budding yeast built on a single CENP-A ortholog (Cse4) nucleosome (Biggins, 2013) to the mega base scale of regional centromeres of humans (Altemose et al., 2022). It has been suggested that regional centromeres could be built from repeated basic subunit that are functionally related to the point centromere. (Allshire & Karpen, 2008). Other centromere types show a similar repeat principle at a larger scale: In the meta-polycentromeric chromosomes of peas (*Pisum sativum*) several discrete CENP-A ortholog (CENH3) domains are distributed along an elongated primary constriction (Neumann et al., 2016). In the holocentromeres of *Rhynchospora pubera* (Hofstatter et al., 2022) many active centromere CENH3 domains are distributed along the entire chromosomes. While many different centromere types have been documented, the basic subunit of centromeres could be similar across these different types and mainly differ in the number and spacing of a common basic subunit, rather than in its nanoscale structure. We hypothesize that if higher-resolution approaches are used to decode centromere architecture in different species, a universal model for centromere architecture maybe identified.

### A multi-domain architecture of the centromere region

Based on published evidence, we previously proposed that human centromeres have a modular, multi-domain architecture (Corless et al., 2025). In our attempt to experimentally validate this model in one single system, our high-resolution imaging revealed three spatially distinct chromatin domains within the centromeric region. The first is the inner kinetochore domain, marked by CENP-A, which occupies the poleward surface of each sister centromere. The second is the outer core domain, marked by CENP-B, which spans the whole primary constriction. The third is the inner centromere domain, marked by RAD21, located between sister chromatids. CENP-B also overlapped with CENP-A, so these domains should be viewed as regions with different protein compositions on a continuous α-satellite array, not as strictly separate compartments.

We could not consistently resolve the proposed mantle domain. RAD21 was detectable next to CENP-A on only 16% of chromosomes. Because this analysis was limited to unexpanded samples, we cannot determine whether this RAD21 lies beneath the CENP-A domain, as the model predicts. The low frequency may reflect a transient or substoichiometric association of cohesin, chromosome-specific differences, or limited antibody detection. Our data therefore neither confirm nor exclude a mantle domain. Imaging and functional studies have suggested that cohesin and condensin contribute to organizing CENP-A chromatin (Sacristan et al., 2024; Sen Gupta et al., 2023). Testing the mantle model directly will require higher-sensitivity detection, for example of endogenously tagged cohesin or condensin subunits at expansion resolution.

Our data confirm at nanoscale resolution that CENP-A occupies only a fraction of the entire centromeric alpha satellite chromatin (Marshall et al., 2008), designated as the inner kinetochore domain or the core centromere as widely referenced to in the literature. This domain serves as the active platform for kinetochore assembly and mediates indirect interactions with spindle microtubules. Our findings underscore a critical point: CENP-A alone does not represent the full centromere region. A comprehensive understanding of centromere function requires conceptualizing it as a multi-domain, modular structure, where each domain contributes distinct functions to centromere integrity and chromosome segregation.

### Histone-modification composition of centromeric and pericentromeric chromatin

Earlier studies of human metaphase chromosomes described centromeres to consist of a distinct chromatin state: CENP-A nucleosomes are interspersed with H3 nucleosomes carrying H3K4me2 and H3K36me2 but lacking H3K9me2/3, while the flanking pericentromeric chromatin carries H3K9me3 and other constitutive heterochromatin marks (Bergmann et al., 2011; Dunleavy et al., 2005; Stellfox et al., 2013; Sullivan & Karpen, 2004). Our expansion microscopy analysis expands this observation to a nanoscale resolution. Across both interphase and metaphase cells, core and pericentromeric chromatin showed strong association with H3S10ph and H3K9me3, low-level association with H3K4me3 and H3K36me3, and lacked of H3K27me3. These observations support the centrochromatin model (Sullivan & Karpen, 2004) in which human centromeric chromatin exists within a spatially complex chromatin environment that includes both canonical heterochromatin and euchromatin associated modifications. However, it needs to be considered that we analysed trimethylated H3K4 and H3K36. Our finding of very low H3K4me3 and H3K36me3 in the CENP-A region is therefore consistent with earlier work but does not directly test the model. Second, the CENP- A region as we defined it may include proximal pericentromeric chromatin at the domain boundaries. The 27% overlap of H3K9me3 with CENP-A may therefore reflect boundary chromatin rather than H3K9me3 interspersed with CENP-A nucleosomes. Our data refine earlier models of pericentromeric chromatin organization (Stellfox et al., 2013) . Rather than being exclusively heterochromatic, the pericentromeric regions analyzed here showed low but reproducible signal for H3K4me3 and H3K36me3. This indicates that open chromatin- associated modifications are present at low abundance within pericentromeric chromatin. However, since we did not resolve individual nucleosomes with our approach, we cannot determine whether euchromatic and heterochromatic marks coexist within the same nucleosome or instead occupy neighboring chromatin subdomains.

The H3S10ph at mitotic chromosomes, including the strong signal observed at centromeric regions fits the known biology of the modification. In G2, H3S10ph first appears at pericentromeric regions and then spreads along the chromosomes (Hayashi-Takanaka et al., 2009). Our observations are consistent with its enrichment in metaphase as well as interphase cells and we observe the strongest H3S10ph signal within centromeric chromatin, including interspersed with CENP-A chromatin. The presence of heterochromatin-associated marks at centromeres and pericentromeres is consistent with their proposed roles in maintaining centromere boundaries and promoting sister-chromatid cohesion (Carty et al., 2025; Mahlke et al., 2026; Sakuno et al., 2009; Yamagishi et al., 2008). The asymmetric H3K9me3 signal observed in some arm-proximal pericentromeric regions further suggests that constitutive heterochromatin may not be uniformly distributed across all pericentromeric domains, although the basis and functional significance of this asymmetry remain to be determined.

H3K27me3 was absent from the centromeric and pericentromeric regions analyzed here. This differs from a report by Lam et al. (2006), who detected low enrichment of H3K27me3 at α-satellite DNA by ChIP-PCR in the HT1080 human cancer cell line. Several factors may explain this difference, including cell-type-specific chromatin states and differences between imaging- and ChIP-based measurements. The low-level detection of H3K4me3 and H3K36me3 in centromeric and pericentromeric regions is consistent with transcription at, which has been implicated in centromere identity or CENP-A deposition (Chueh et al., 2009).

Overall, our expansion microscopy analysis shows that mitotic centromeric and pericentromeric chromatin is dominated by H3S10ph and H3K9me3, lacks H3K27me3, and has only low levels of H3K4me3 and H3K36me3. Future studies combining nucleosome- resolution imaging, chromatin profiling, and functional perturbation will be required to determine how these histone modifications contribute to centromere identity, cohesion, and kinetochore function.

## Limitations

This study relies primarily on expansion microscopy to visualize centromeres at nanoscale resolution. Although this approach improves resolution using conventional microscopes, it depends on antibody specificity, epitope accessibility and labelling efficiency. Thus, despite using validated antibodies, these factors may influence the accuracy of our observations. In addition, low-abundant targets are more difficult to detect by increasing the distance between fluorophores after physical expansion. In addition, although our protocol was designed for 10× expansion, we typically achieved expansion factors of 7×, likely reflecting variability in sodium acrylate quality. Because this variability can affect absolute distance measurements, we did not report extensive quantitative distance-based analyses. In addition, we relied on imaging one cell line, hTERT RPE-1. Centromere size and α-satellite organization differ between chromosomes and between individuals, so the range of CENP-A subdomains may vary in other cell types. We also could not trace the identity of chromosomes in our images. As a result, we could not compare subdomain counts for specific centromeres directly with sequencing-based sub-CDR counts, and the relationship between imaged CENP-A foci and sub-CDRs remains correlative.

## Materials and methods

### Antibodies

| Target | Supplier | Catalogue number |
|---|---|---|
| CENP-A | Enzo Life Sciences | ADI-KAM-CC006-E |
| KNL1 | Sigma-Aldrich | HPA026624 |
| CENP-B | Abcam | ab25734 |
| RAD21 | Abcam | ab154769 |
| H3K4me3 | Abcam | ab8580 |
| H3K36me3 | Abcam | ab9050 |
| H3K9me3 | Abcam | ab8898 |
| H3K27me3 | Cell Signaling Technology | 9733 |
| H3S10ph | Abcam | ab5176 |
| Anti-rabbit Alexa Fluor 488 | Thermo Fisher Scientific | A11008 |
| Anti-mouse Alexa Fluor 594 | Cell Signaling Technology | 8890S |

### Cell culture and immunofluorescence

hTERT RPE-1 cells were cultured in DMEM/F-12 supplemented with 10% fetal bovine serum, 1% penicillin-streptomycin, 2 mM L-glutamine and 0.348% sodium bicarbonate at 37 °C, 5% CO₂. Cells were seeded on 12-mm glass coverslips in 24-well plates. Mitotic cells were enriched by treatment with colcemid at 100 ng/mL concentration for 4-6 h. For immunofluorescence, cells were pre-extracted for 1 min at room temperature in extraction buffer containing 60 mM PIPES, 25 mM HEPES, 2 mM MgCl₂·6H₂O, 10 mM EGTA and 0.5% Triton X-100. Cells were then fixed in 4% formaldehyde for 10 min at room temperature and blocked in 5% BSA overnight. Primary antibodies were diluted 1:100 in blocking buffer and incubated for 3 h at room temperature. Secondary antibodies were diluted 1:100 and incubated for 1 h at room temperature. 0.5% PBST was used as the buffer for incubation and washes throughout. For RAD21 staining, all incubations and washes were performed in KCM buffer containing 120 mM KCl, 20 mM NaCl, 10 mM Tris-HCl pH 8.0, 0.5 mM EDTA and 0.1% Triton X-100.

### Metaphase spreads and immunofluorescence

Following colcemid treatment, cells were incubated in hypotonic 0.075 M KCl for 10 min at room temperature. Approximately 3 × 10⁴ cells were cytospun onto coverslips using a Thermo Scientific Cytospin 4 at 1800 rpm for 10 min using the low-acceleration setting. Immunofluorescence was then performed as described above.

### Ten-fold robust expansion microscopy

Ten-fold robust expansion microscopy was performed as described by Damstra et al. (2023). Briefly, after immunofluorescence, coverslips were incubated in anchoring solution containing 3% acrylamide and 1.2% formaldehyde for 3 h at 37 °C. Coverslips were then placed cell-side down on gelation solution containing 1.1 M sodium acrylate, 14.4% acrylamide, 0.005% N,Nʹ- methylenebisacrylamide, 0.15% TEMED, 0.15% APS and 1× PBS. Gels were polymerized for 1.5 h at 37 °C. Samples were digested for 4 h at 37 °C in digestion buffer containing 50 mM Tris- HCl pH 8.0, 1 mM EDTA, 0.8 M guanidine hydrochloride and 0.5% Triton X-100 supplemented with 200 µg/mL proteinase K (Sigma 1.24568). Gels were expanded by repeated exchange into water every 30 min until fully expanded.

### Mounting and microscopy

Conventional immunofluorescence samples were mounted in Vectashield containing DAPI. For expansion microscopy, gels were trimmed and mounted on poly-L-lysine-coated coverslips in an Attofluor cell chamber A7816. Images were acquired on a Zeiss Airyscan LSM 900 microscope using Plan-Apochromat 63×/1.4 NA oil DIC M27 objective. Images were acquired using Airyscan multiplex 4Y mode. Airyscan joint deconvolution or two-dimensional deconvolution was performed after acquisition using ZEN 3.13.

### Image analysis

Image analysis was performed in Huygens Essential v25.10. Chromatic aberration was corrected using multicolor bead images taken using the same microscope settings as a reference and the Chromatic Aberration Chromatic wizard. Three-dimensional rendering was performed using the Object Analyzer module. Signals were segmented using the watershed algorithm with default parameters. Three-dimensional colocalization between CENP-A and the indicated histone marks and RAD21 was quantified using the colocalization (intersection) function in Object Analyzer. Data were exported as CSV files and analyzed in RStudio. Box plots show relative colocalized volume (relColocV %), defined as the percentage of CENP-A voxels intersecting with histone mark/RAD21 positive voxels in 3D. To quantify CENP-A domain number, line-scan intensity profiles were manually generated, and individual peaks with fluorescence intensities above 1,000 arbitrary units (a.u.) were scored as distinct CENP-A domains.

## Data availability

The metadata associated with the microscopic images are available upon request.

## Author contributions

Conceptualization: GT, SE

Methodology & investigation: GT

Funding acquisition: SE

Writing, review and editing: GT, SE

## Acknowledgments

We thank Franco Weth, Mark E. Pownall, Anneloes Keijzer, Carlos Sacristan Lopez, and Geert Kops for sharing their expertise in expansion microscopy. We also thank the members of the KIT VirtMat team, the SynGen team, and the Erhardt laboratory for helpful discussions throughout the project. Figure 4 was created using BioRender. We acknowledge funding from Synthetic Genomics (SynGen) and the Carl Zeiss Stiftung.

**Figure 4.**
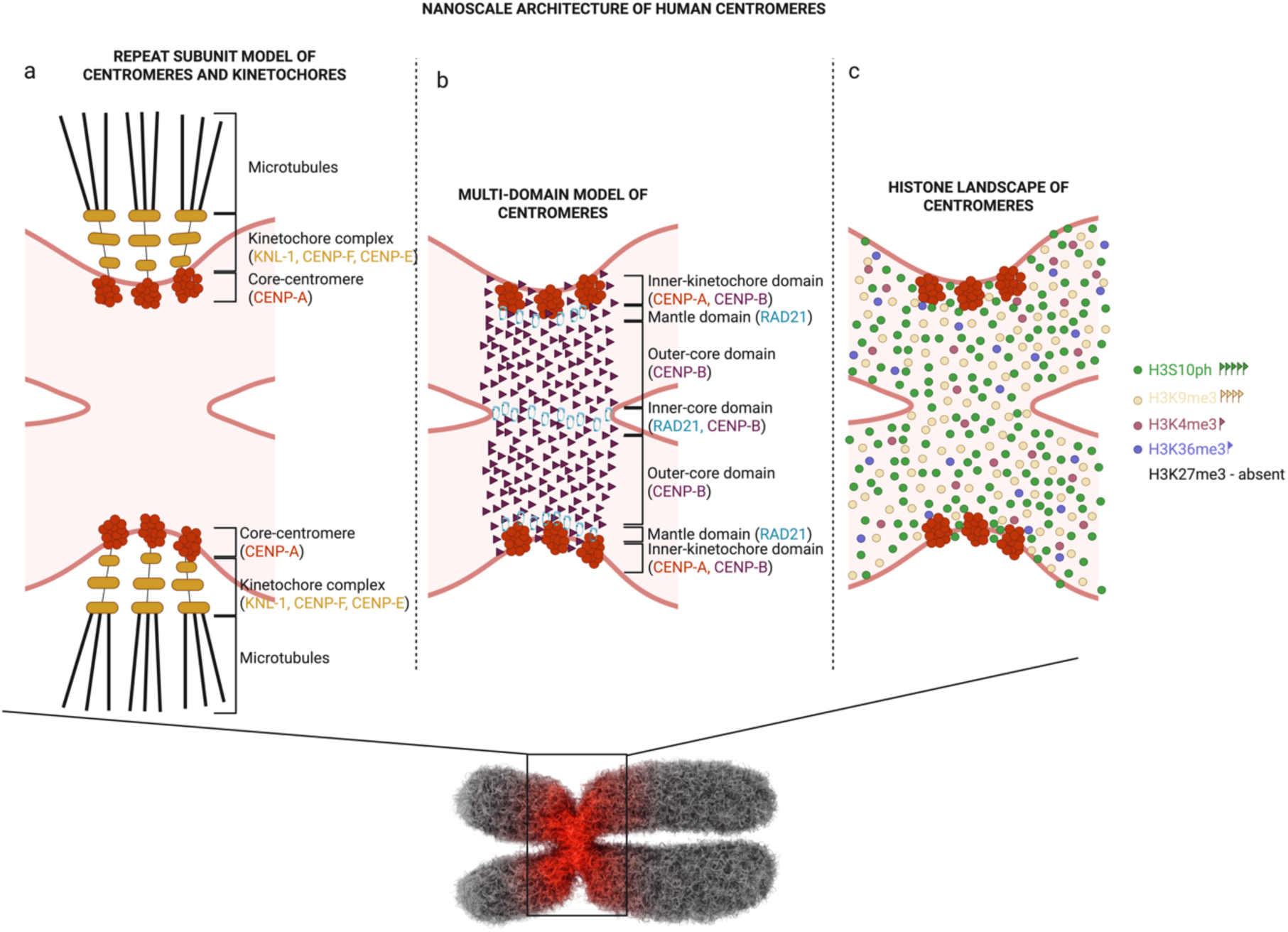
Model for the architecture of human centromeres. **a**, Multi-domain model of centromere organization, showing four distinct domains (inner kinetochore, mantle, outer core, and inner core) that together define the centromere architecture. **b,** Repeat subunit model of centromeres and kinetochores, illustrating that multiple discontinuous subunits form the underlying three-dimensional organization of centromeres and kinetochores. **c,** Histone landscape of centromeres, depicting the spatial distribution of different histone modifications across centromeric chromatin.

---

**Supplementary Fig. 1.**
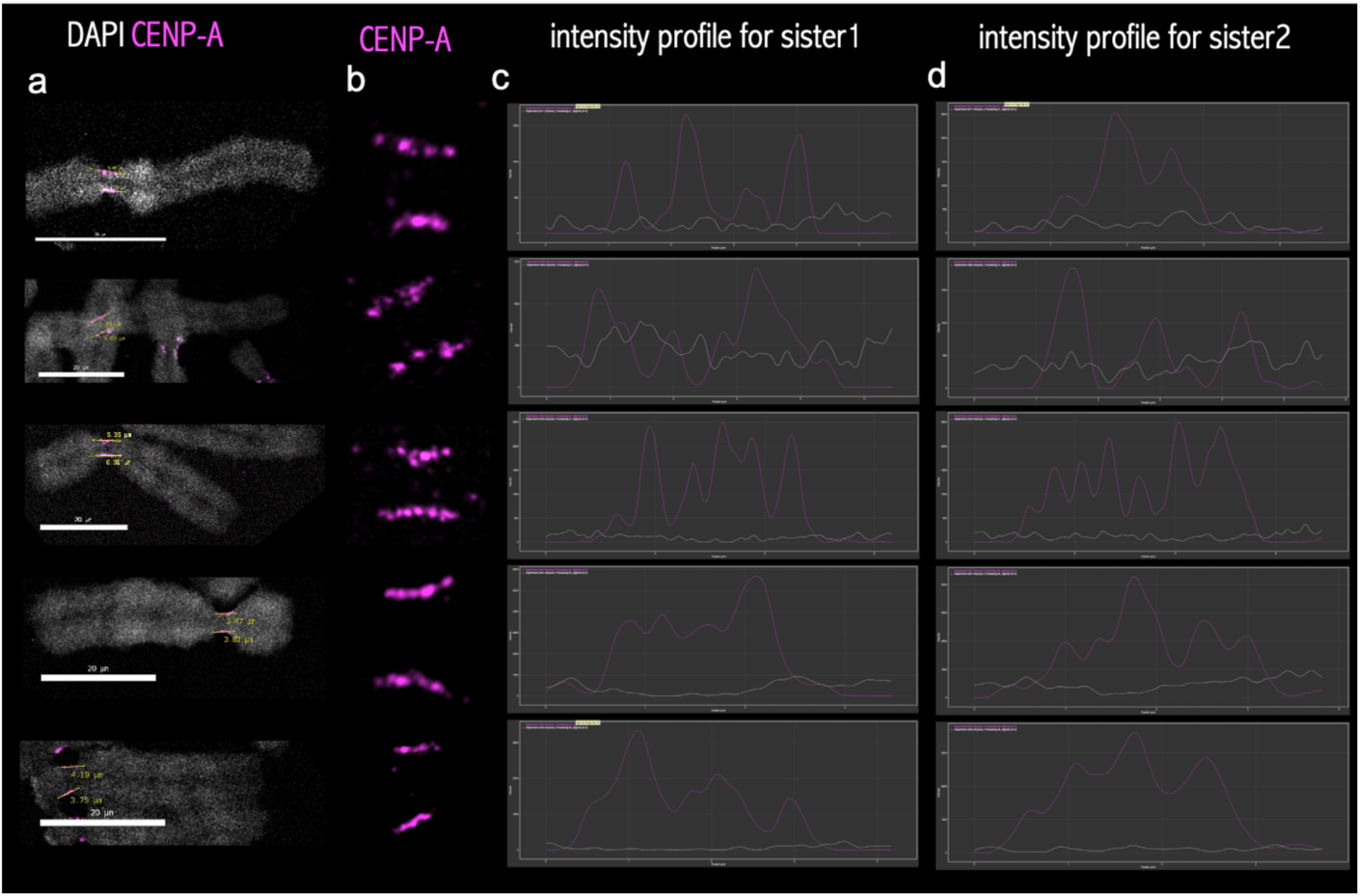
Multiple discontinuous CENP-A subunits revealed by expansion microscopy. **a,b**, Representative chromosomes stained for CENP-A (magenta), showing multiple discontinuous subunits. **c,d,** Fluorescence intensity profiles across sister chromatids, revealing multiple CENP-A peaks within each sister chromatid. Scale bar, 20 μm.

**Supplementary Fig. 2.**
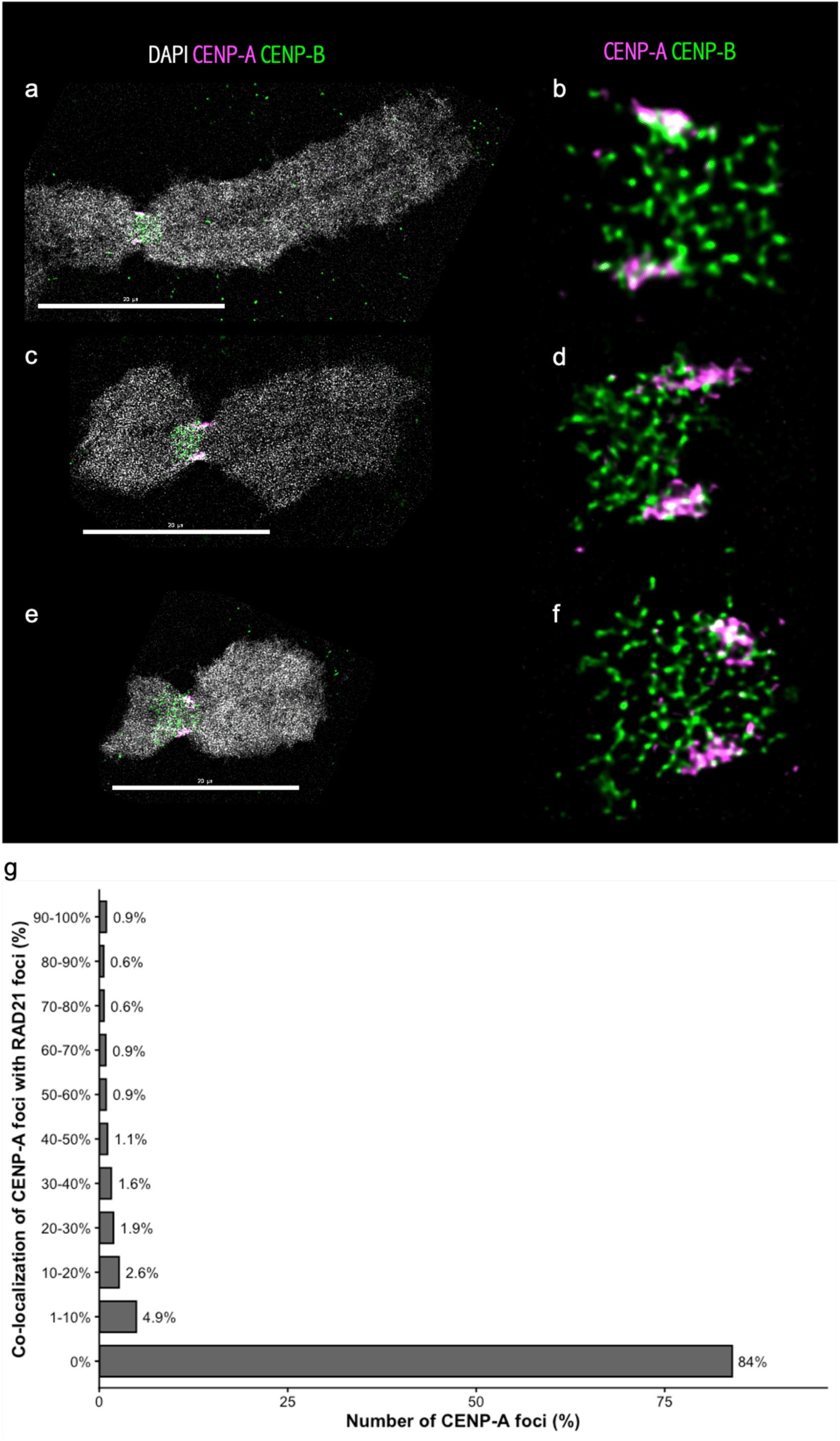
CENP-B distribution on metaphase chromosome spreads. **a–f**, Representative images of metaphase chromosomes stained for CENP-A (magenta) and CENP- B (green), showing asymmetric CENP-B distribution between chromosome arms. One chromosome arm displays stronger CENP-B enrichment compared with the opposite arm. **g,** distribution of co-localization values in % between CENP-A and RAD21. Scale bar, 20 μm.

**Supplementary Fig. 3.**
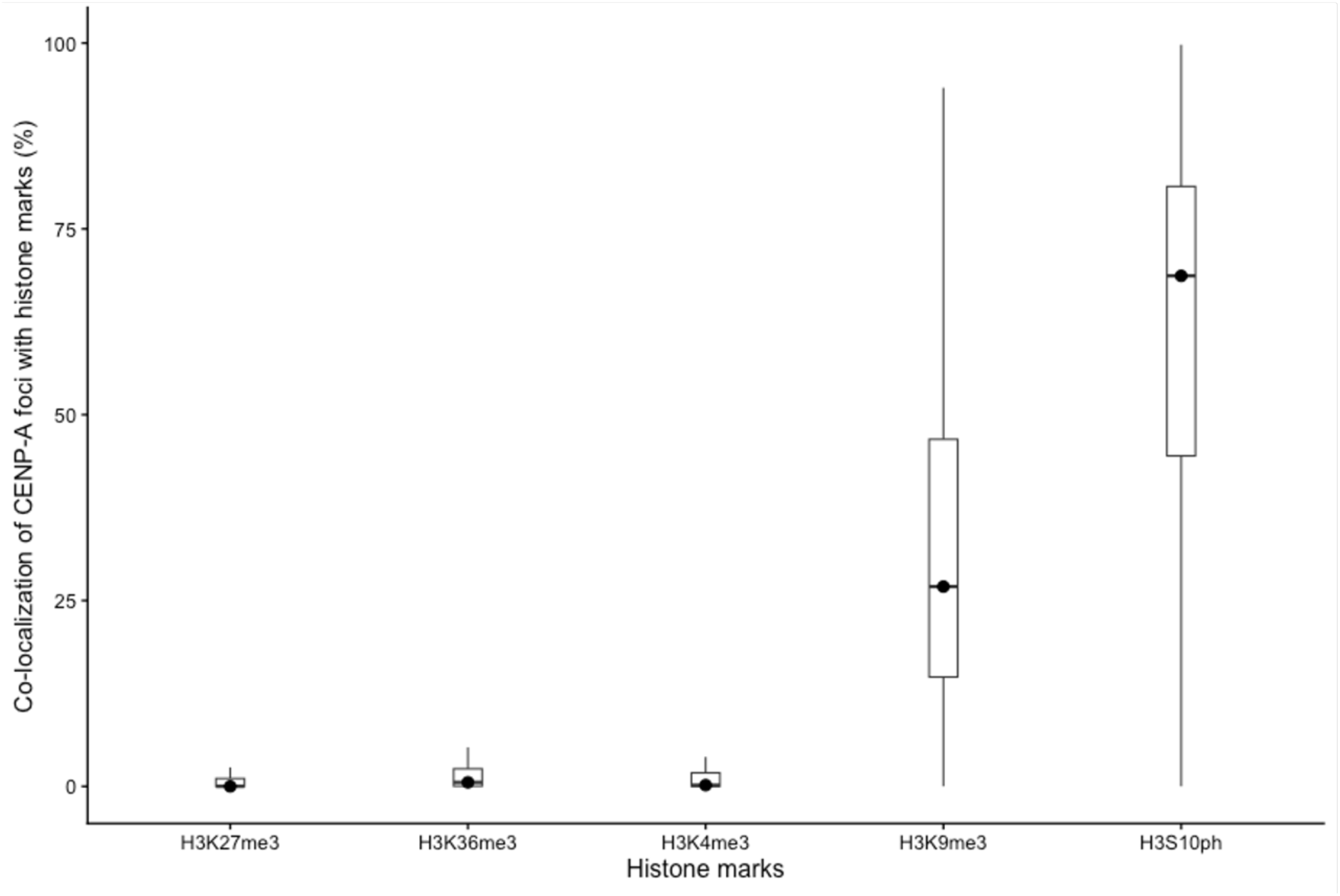
Three-dimensional co-localization analysis of CENP-A and different histone modifications. The box plot shows the percentage of CENP-A (measured as voxels in 3D) that co-localizes with different histone marks H3K27me3, H3K36me3, H3K4me3, H3K9me3 and H3S10ph during metaphase.

